# Evolution of nuptial gifts and its coevolutionary dynamics with “masculine” female traits for multiple mating

**DOI:** 10.1101/2020.06.22.165837

**Authors:** Yoshitaka Kamimura, Kazunori Yoshizawa, Charles Lienhard, Rodrigo L. Ferreira, Jun Abe

## Abstract

Many male animals donate nutritive materials during courtship or mating to their female mates. Donation of large-sized gifts, though costly to prepare, can result in increased sperm transfer during mating and delayed remating of the females, resulting in a higher paternity Nuptial gifting sometimes causes severe female-female competition for obtaining gifts (i.e., sex-role reversal in mate competition) and female polyandry, changing the intensity of sperm competition and the resultant paternity gains. We built a theoretical model to analyze such coevolutionary feedbacks between nuptial gift size (male trait) and propensity for multiple mating (female trait). Our genetically explicit, individual-based computer simulations demonstrate that a positive correlation between donated gift size and the resultant paternity gain is a requisite for the co-occurrence of large-sized gifts and females’ competitive multiple mating for the gifts. When donation of gifts imposes monandry, exaggeration in nuptial gift size also occurs under the assumption that the last male monopolizes paternity. We also analyzed the causes and consequences of the evolution of a female persistence trait in trading of nuptial gifts, that is, double receptacles for nuptial gifts known to occur in an insect group with a “female penis” (*Neotrogla* spp.).

## 1. INTRODUCTION

Nuptial gifts, any non-gametic materials transferred from one sex (usually male) to another during courtship and mating, are widely observed in many groups of animals, such as insects, arachnids, molluscans, amphibians, birds, and mammals including humans (Lewis and South 2012; Lewis et al. 2014). In some cases, male-derived “gifts” can be detrimental to female recipients as the love darts of land snails and anti-aphrodisiac seminal peptides of *Drosophila* fruit flies, both of which mitigate the intensity of sperm competition in female sperm storage organs (Chapman 2001; Chase and Blanchard 2006). However, in many cases, males donate (at least potentially) nutritious materials such as prey items they have collected or voluminous secretions from male internal/external glands (Boggs 1995; Lewis and South 2012).

Although this phenomenon is widely observed in the animal kingdom, there is continuing debate on its primary function (Alexander and Borgia 1979; Gwynne 1984; Sakaliuk 1986; Simmons and Parker 1989; Parker and Simmons 1989; Wedell 1993; Wickler 1994; Vahed 1998). Since females of some animals accept mating only while consuming a gift, males may donate the gift to obtain mating opportunities (i.e., mating effort hypothesis). In addition to nourishing female recipients, nutrition from nuptial gifts can be passed to offspring sired by the male donor, and thus can function as paternal investments (i.e., paternal investment hypothesis). Regardless of the ultimate benefits for males, a large nutritious gift is costly for the male to prepare (reviewed in Boggs 1995), although it may be more attractive to females or may enable transfer of more sperm (Sauer et al. 1998; Engqvist et al. 2007; South and Lewis 2012). By improving the nutritional status of female recipients, gigantic nuptial gifts can thus also be a cause of reversal in the sex roles: usually males more actively seek mating opportunities while females are choosier about mates than males, but in sex-role reversed animals, females compete for multiple mating opportunities to obtain more nuptial gifts (Vahed 1998; Gwynne 2008; Fritzsche & Arnqvist 2013; Kamimura and Yoshizawa 2017). Increased polyandry inevitably causes severer sperm competition, changing the cost-benefit balance for males to prepare nuptial gifts. Although several models have been proposed to date for elucidating the evolution of nuptial gifts (Parker and Simmons 1989; Boggs 1990; Alonzo and Pizzati 2010), they did not encompass these possible coevolutionary feedbacks between the sexes for the trading of gifts. The patterns of female sperm storage and use, which determine the benefits in paternity gain of donating a given gift size of gifts, are possible pivotal factors for shaping the coevolutionary dynamics in the trading of nuptial gifts. However, explicit appraisals, both theoretical and empirical, are lacking entirely.

To fill this gap, we developed a genetically explicit individual-based model in the present study. For this purpose, we chose *Neotrogla* (Insecta: Psocodea: Prinoglarididae: Sensitibillini) as an illustrative case. A single male ejaculate of *Neotrogla*, which contains voluminous and potentially nutritional seminal substances, forms a gigantic, bottle-shaped capsule (spermatophore) of ~0.05 mm^3^, corresponding to ~300 ml if scaled up to human proportions (Yoshizawa et al. 2014, 2018b). Unlike prey items, the size of which cannot be exactly controlled by the male donors, males likely allocate their limited resources for this type of seminal gift to multiple mating opportunities in a strategic manner. This model system provides us the rare opportunity to examine additional complexities in the evolution of sex-role reversed animals, that is, the evolution of “masculine” persistence traits in females for competitively obtaining male-derived gifts. Females of *Neotrogla* use their evolutionarily novel penis-like structure, termed a gynosome, as an intromittent organ for copulation with males. This “female penis” is ornamented with species-specific lobes and/or spine bundles, which are accommodated in specialized pouches of the vagina-like male genitalia during copulation (Yoshizawa et al. 2014). Since they live in dry, nutritionally poor caves in Brazil, the gynosome likely represents an elaborate way for females to exploit seminal gifts from males (Yoshizawa et al. 2014, 2019a). In accordance with this view, females of a related species with similar spermatophores (Psocodea: Trogiidae: *Lepinotus*) compete for access to males (Wearing-Wilde 1995, 1996). Moreover, females of *Neotrogla* and those of related genera of the tribe Sensitibillini (*Afrotrogla* and *Sensitibilla*) have also developed a specialized structure, termed a spermathecal plate, in their sperm storage organ (spermatheca). In *Neotrogla* (and possibly also in *Afrotrogla* and *Sensitibilla*), this evolutionarily novel organ is equipped with twin slots that enable retention and digestion of two seminal gifts simultaneously (Lienhard 2007; Yoshizawa et al. 2014, 2018b, 2019a). By contrast, females of related groups with only a single slot for nuptial gift can accept another mating only after digestion of the content of a spermatophore received at the preceding mating (for example, in *Lepinotus*, Wearing-Wilde 1995). Given that *Neotrogla* females mate multiply as evidenced by up to two full and nine emptied spermatophore capsules in their spermatheca (Yoshizawa et al. 2014), the spermathecal plate of this genus (and probably also of the remaining Sensitibillini) can be considered a female adaptation for obtaining nutritious gifts in rapid succession.

In the present study, we examine coevolutionary dynamics between the size (volume) of nutritive seminal gifts, a male trait, and propensity for multiple mating to obtain gifts, a female trait. Resource availability to males, costs of additional mating for females, and the paternity determination patterns in multiply mated females were analyzed as possible factors influencing the coevolutionary dynamics. The evolutionary causes and consequences of females with twin slots for obtaining gifts, hereafter referred to as “*2S* females”, were also examined in detail to understand the evolution of “masculine” traits in sex-role reversed female animals.

## 2. MODEL ASSUMPTIONS

All notations and parameter values used in this article are summarized in Table 1. It is likely that a given volume of nuptial gift will increase the female’s fitness more effectively when the female is starving than when she has already received a large amount of nutrients from the preceding mates. Thus, we assumed that the potential offspring number of a female (fecundity, *F_pot_*) is a saturation function of the cumulative volume of seminal gifts (*r*) received in all of her previous matings (Fig. 1):

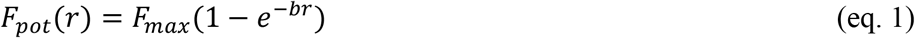

where *b*, set at 0.002 throughout the present study, represents the speed of saturation. In this equation, *F_max_*, set at 800, denotes the maximum number of offspring that a female can potentially produce when *r* = ∞ under no cost of mating.

**Fig. 1.**
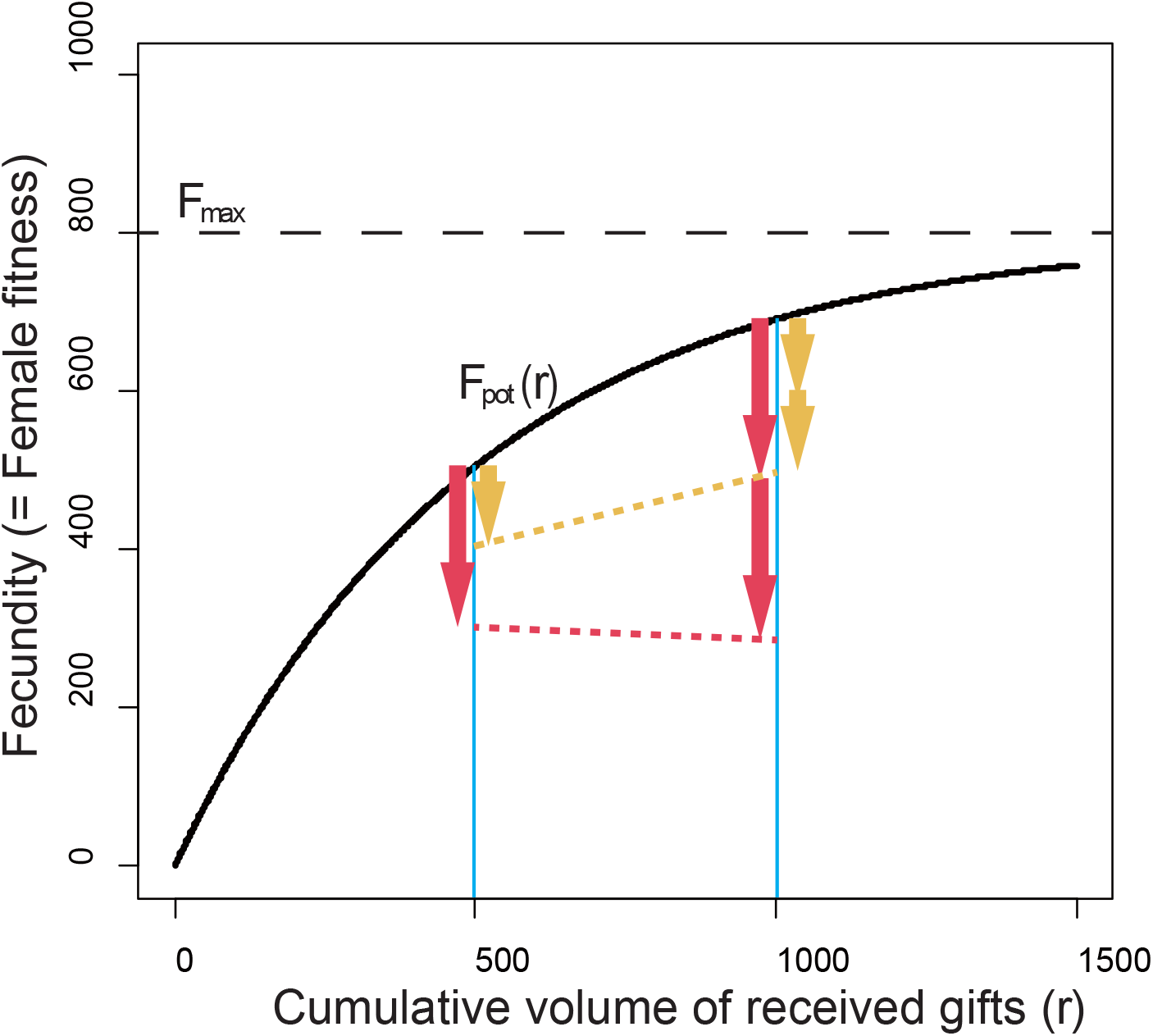
Female fecundity as a saturating function of seminal nutrients from males. See eqs. (1) and (2) in the main text. When males provide a seminal gift of size 500 (*V* = 500), females that mated once should seek another mating opportunity when the cost of mating is low (orange arrows, *c* = 100), but not under a higher cost (red arrows, *c* = 200).

**Table 1.** Summary of notation and abbreviations used in this study.

| Symbol | Definition and designated values |
| --- | --- |
| **MODEL PARAMETERS** |  |
| $v1, v2$ | Genetical values of male seminal gift size from male and female parents, respectively. Initial values were given as a normal distribution (mean ± SD = $20 \pm 40$) with truncation ($\geq 20$). |
| $V$ | Phenotypic value of male seminal gift size, calculated as $(v1 + v2)/2$. |
| $V_{last}$ | Seminal volume of the last mating of a focal male, given as eq. 3 |
| $m1, m2$ | Genetical values of female acceptable number of mating from male and female parents, respectively. Initial values were given as a normal distribution (mean ± SD = $0.5 \pm 2$) with truncation ($\geq 0$). |
| $M$ | Phenotypic value of female acceptable number of additional mating (= propensity for multiple mating), calculated as $(m1 + m2)/2$. |
| $M_R$ | Realized number of mating by females |
| $M_O$ | Optimal number of mating by females calculated by solving eq. 2 |
| $2s$ | Dominant gene for producing twin slots for receiving gifts |
| $1s$ | Recessive gene for producing a single slot for receiving gifts |
| $2S$ | Females of the genotype $2s/2s$ or $2s/1s$, having twin slots for insemination |
| $1S$ | Females of the genotype $1s/1s$, having a single slot for insemination |
| $R$ | Average total volume of resource for producing seminal gifts, ranged 400-1300 by an increment of 100 depending on the resource availability of the habitat. Each male has a resource budget extracted from a truncated normal distribution mean ± SD = $R \pm 0.2R$ ($> 0$). |
| $N$ | Possible number of mating for a focal male, given as $R/V$, rounded up to the nearest integer. |
| $r$ | Cummulative volume of gifts received by a female |
| $b$ | Fertilization efficiency (this parameter represents the saturating speed of female reproductive output). Treated as a constant ($b = 0.002$) in this study. |
| $F_{max}$ | Maximum female reproductive output (e.g., offspring number of a female when $r = \infty$ and $c = 0$). Treated as a constant ($F_{max} = 800$) in this study. |
| $c$ | Mating cost for females, ranged 5-95 by an increment of 10 |
| $F_{pot}(r)$ | Maximum fecundity of a female, as a function of $b, k$ and $r$, before reduction by mating costs |
| $F(r, c, M_R)$ | Realized fecundity of a female, as a function of $b, k, r, M_R$, and $c$. |
| **PATERNITY DETERMINATION REGIMES** |  |
| PC | Paternity is perfectly correlated with the relative gift size |
| ES | Paternity is equally shared by all male mates |
| LM | The last male monopolizes paternity |
| FM | The first male monopolizes paternity |
| **SIMULATION MODES** |  |
| CON | The control simulation runs with neither invasion of $2S$ females nor doubling in female number |
| DS | One $2S$ female per generation was introduced from 1000th generation and beyond |
| DF | The number of females were doubled at 1000th generation (i.e., 1000 females for 500 males) |

Even in cases in which males donate nuptial gifts at each mating event, multiple mating, which may involve costly mate-searching and an enhanced risk of being predated, can be detrimental for the females. Thus, we imposed a cost (*c*) for each mating event, as a reduction in the number of offspring as:

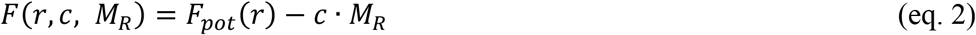

where *M_R_* is the number of performed matings of a focal female.

This simple function delineates complicated coevolutionary relationships between the male and female traits (Fig. 1). When males donate a large-sized gift at each mating (e.g., *V* = 500), an additional mating further increases the lifetime fitness of a singly-mated female under a low cost mating (e.g., *c* = 100; orange arrows in Fig. 1), but not when it is largely costly (e.g., *c* = 200; red arrows). From a male perspective, because the fitness gain to a female mate from receiving a unit amount of gift gradually decreases with the total amount of gifts received in previous rounds of mating (*r*, Fig. 1), preparing small-sized gifts is advantageous because it increases opportunities for mating with females, especially unmated females, who received less or no nutrition from previous matings. However, smaller gifts can result in a higher probability of remating by his female mates when mating cost is sufficiently low. Therefore, males who donate a large-sized gift can enhance the paternity gain by decreasing the probability of remating by their female mates, although the large-sized gift inevitably decreases the number of matings in which that they can engage. Furthermore, in the case that males can transfer more sperm by giving a large-sized gift, it can result in a higher paternity share compared to males who gave a smaller gift to the same mate. Since sperm storage/use patterns of multiply-mated females are largely unknown for animals with donation of nuptial gifts (Lewis and South 2012), we examined four different regimes for the relationship between the nuptial gift size and the resultant paternity gain in this study: (1) perfect correlation (PC), (2) equal shares (ES), (3) complete last male (LM), and (4) complete first male (FS) (Fig. 2, Table 1).

**Fig. 2.**
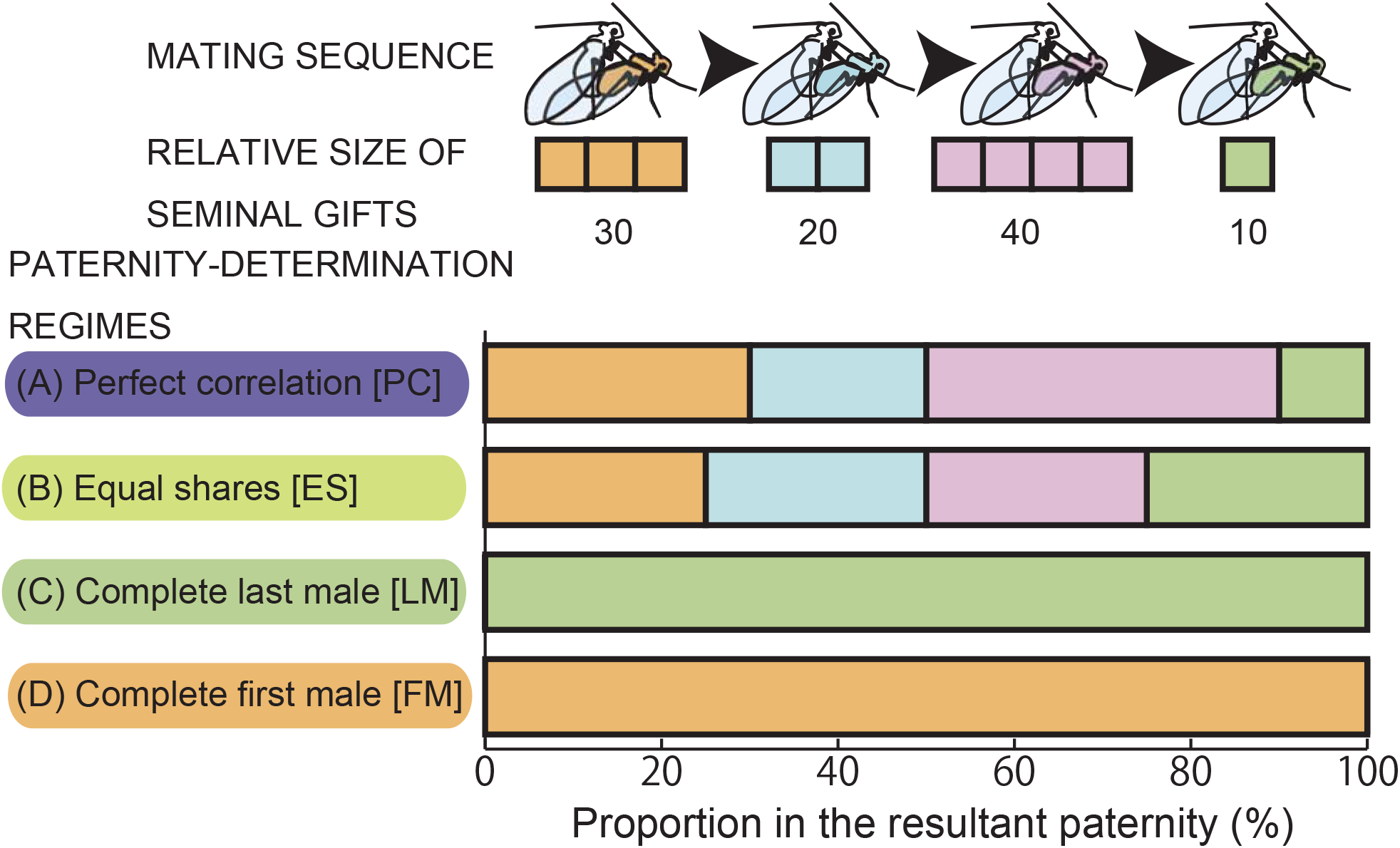
A scheme of four different regimes of the relationships between seminal gift size and the resultant paternity. Note that *Neotrogla* shows a female-above mating posture similar to many other species of Psocodea.

## 3. INDIVIDUAL-BASED SIMULATIONS

Individual-based evolutionary simulations were conducted to observe the coevolutionary dynamics of three traits: male seminal gift size, female propensity for multiple mating, and the number of slots for obtaining gifts in females. For this, we used personal scripts written in Python 3.7.1, which are available in the Appendix. Simulations were run for 2000 generations assuming a single population of diploid organisms. The population size was set at 1000 (500 males and 500 females). Nuptial gift size, a trait of male-specific expression, was assumed to be determined by a set of autosomal polygenes. To mimic sexual reproduction, these polygenes were divided into two groups, one from each parent. The initial genetic value for each polygene group was randomly extracted from a normal distribution, with a mean of 20 and a standard deviation (SD) of 40. Each individual possesses two values, *v1* and *v2*, as the genotype of this trait. The seminal gift size (volume; *V*) was determined as the average of these two values only for male individuals as their phenotype. Similarly, a mean of 0.5 ± 2 was given as the initial values for the propensity for multiple mating (*m1* and *m2*), which determines the maximum number of “additional” matings accepted by each female individual (*M*), as the nearest integer of (*m1* + *m2*)/2. Thus, females of the genotype *m1/m2* mate *M* + 1 times, whenever a mating opportunity is available. To prevent the occurrence of unreasonable values in these two traits, we set the lower limits of genetic values of these two traits as 20 and 0, respectively.

Another female trait, the number of slots for accepting nuptial gifts, was assumed to be a dichotomous trait, that is, one or two slots. Little is known about the evolutionary process of the spermathecal plate, which enables retention of two nuptial gifts simultaneously, in the ancestors of extant Sensitibillini (see Discussion). For simplicity, we assumed that the transition is controlled by a single locus with two alleles: *2s*, a gene for the twin-slot state, and *1s*, a gene for the single-slot state, the former was assumed to be dominant over the latter. The populations were initiated with single-slot females (*1S*, *1s*/*1s* homozygotes). Then, *2s* genes were allowed to invade to populations at 1000th generation and beyond (one spontaneous mutation in each generation), together with recurrent invasion of single-slot mutants (a female individual of *1s*/*1s* homozygote per generation) to check the evolutionary stability of the twin-slot state (*2S*). When the *2S* phenotype occupied 95% or more of females, this phenotype was judged as fixed.

In the model, each male can mate up to *N* times, which is *R/V* rounded up to the nearest integer, where *R* is a total volume of resource available for production of nuptial gifts, randomly extracted from a normal distribution. The gift size of the last mating (*V_last_*) is:

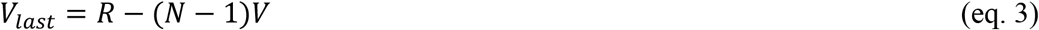

unless *R* is not a multiple of *V*.

The mean value of *R* ranged from 400 to 1300, by increments of 100, representing an environmental variability in resource availability. The SD was set at each mean value multiplied by 0.2.

In our model, a female accepts additional mating when (1) she has mated fewer times than her acceptable number of matings (*M* + 1), and (2) at least one of her slots is empty. We assumed that in every generation, all virgin females start to mate simultaneously. After the first mating between random male and female pairs, seminal gifts from males were sequentially assigned, one at a time, to the slot that had received the minimum cumulative volume of gifts (*r*) in the females whose mating demand was not satisfied at the time. This procedure was repeated until all females satisfied their mating demands, or all males exhausted their resource budgets. By this algorithm, we implemented the condition that larger seminal gifts are difficult to digest for females, and thus occupy a slot for a longer duration, resulting in delayed remating of the female.

Then, the relative fitness of each female was calculated according to eq. 2. Cost of mating, defined by a reduction in female fitness per mating (Fig. 1), was varied from 5 to 95, by increments of 10, and negative fitness values were treated as zero. Maternity and paternity of offspring were determined according to the proportional representation of the relative fitness of females and the relative representation of male sperm in each female (Fig. 2), respectively. For each trait, one of two parental genes was randomly and independently extracted from two parents, and fused to create an individual of the next generation, that is, with no linkage among the three traits. For seminal gift size and propensity for female multiple mating, each of these values was treated as the mean of a normal distribution for creating the genetic values of progenies (recurrent mutation) with SDs 40 or 2, respectively.

To dissect the complicated coevolutionary interactions between the male and female traits, simulations were repeated 40 times for each combination of *R* and *c* values, and for each of the four different paternity determination regimes (PC, ES, LM, and FM) specified above (Fig. 2). In addition, three different types of simulations were conducted (Table 1). In doubling females (DF) runs, the number of females was doubled, resulting in 1000 females per 500 males, from 1000th generation and beyond, instead of introducing invasion of one twin-slot mutant female (*2S* female) every generation (doubling slots, DS runs). In the control runs (CON), neither the numbers of slots nor females were doubled throughout the 2000-generation runs. Possible effects of genetic correlation between the male and female traits on their coevolution were also examined (see Supporting Material 2).

## 4. RESULTS

### Evolution of nuptial gift size and female multiple mating under different paternity-determination regimes

Our simulations revealed notable effects of the pattern of paternity determination on the evolution of gigantic seminal gifts and female persistence traits, that is, the propensity for multiple mating and twin slots for obtaining gifts. Under a given set of parameters, the male seminal gift size and female propensity for multiple mating rapidly converged to a respective equilibrium, usually before 300th generation (the left half of Fig. 3 for an example of DS runs; for examples of CON and DF runs, see Supporting Material 1). Only when females used sperm from each male mate for fertilization of eggs proportionally to the nuptial gift size donated (the PC regime) did males evolve a large-sized gift compared with their lifetime resource budget under a wide range of parameter sets (male resource budget [*R*] and cost of mating for females [*c*]: Fig. 4A).

**Fig. 3.**
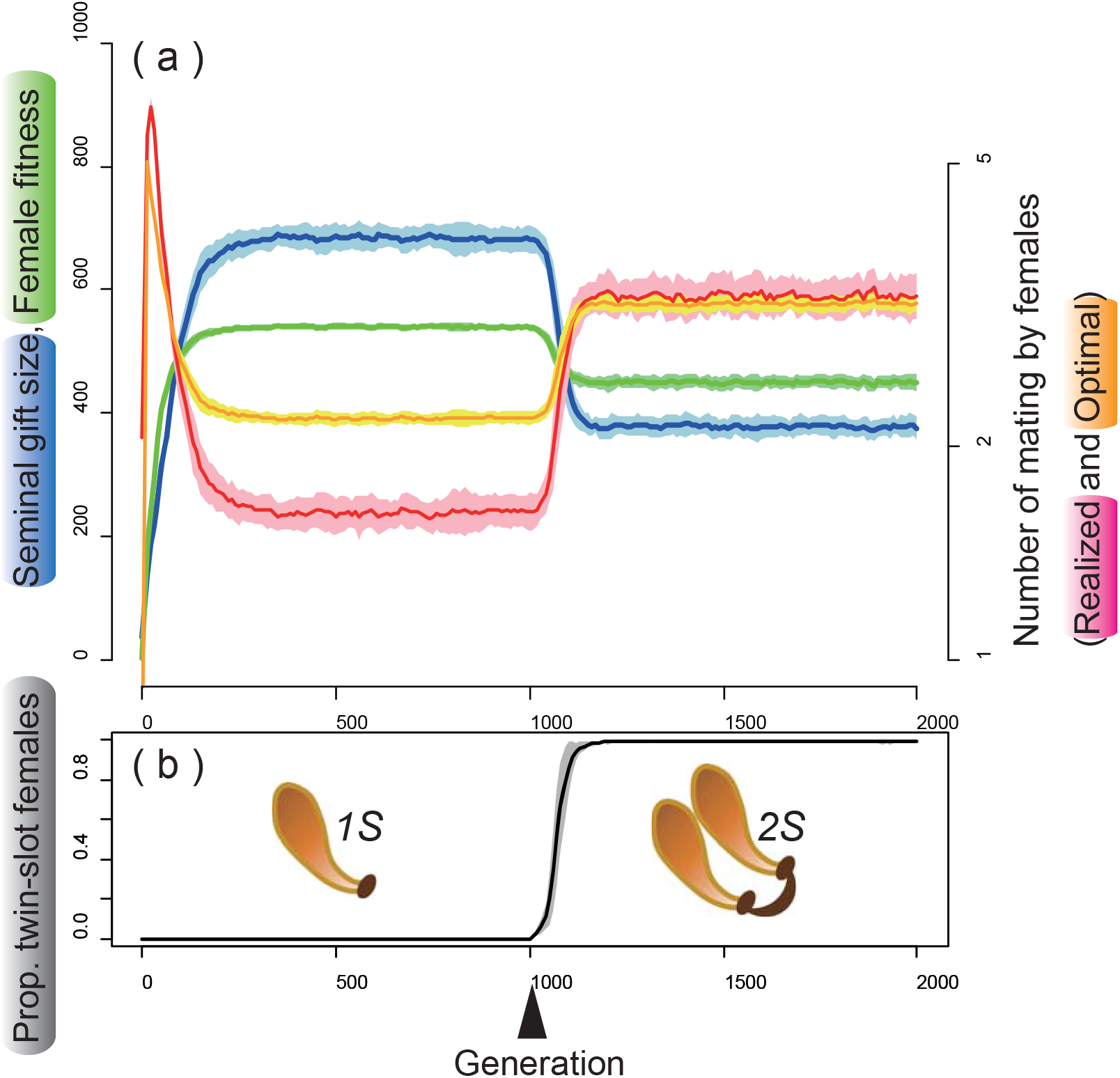
**(a)** An example of the coevolutionary dynamics observed between the male seminal gift size (*V*, blue) and the optimal number of matings for females (*M_o_*, red), together with changes in the realized number of matings by females (*M_R_*, orange) and female fitness (*F*, green). **(b)** Changes in the proportion of females with twin slots (*2S*), which started to invade the population at the 1000th generation (the black arrowhead), with schematics of *1S* and *2S* states. Solid lines and shaded areas of respective lighter colors show the mean ± SD for 40 runs under the PC regime (*R* = 800, *c* = 55).

**Fig. 4.**
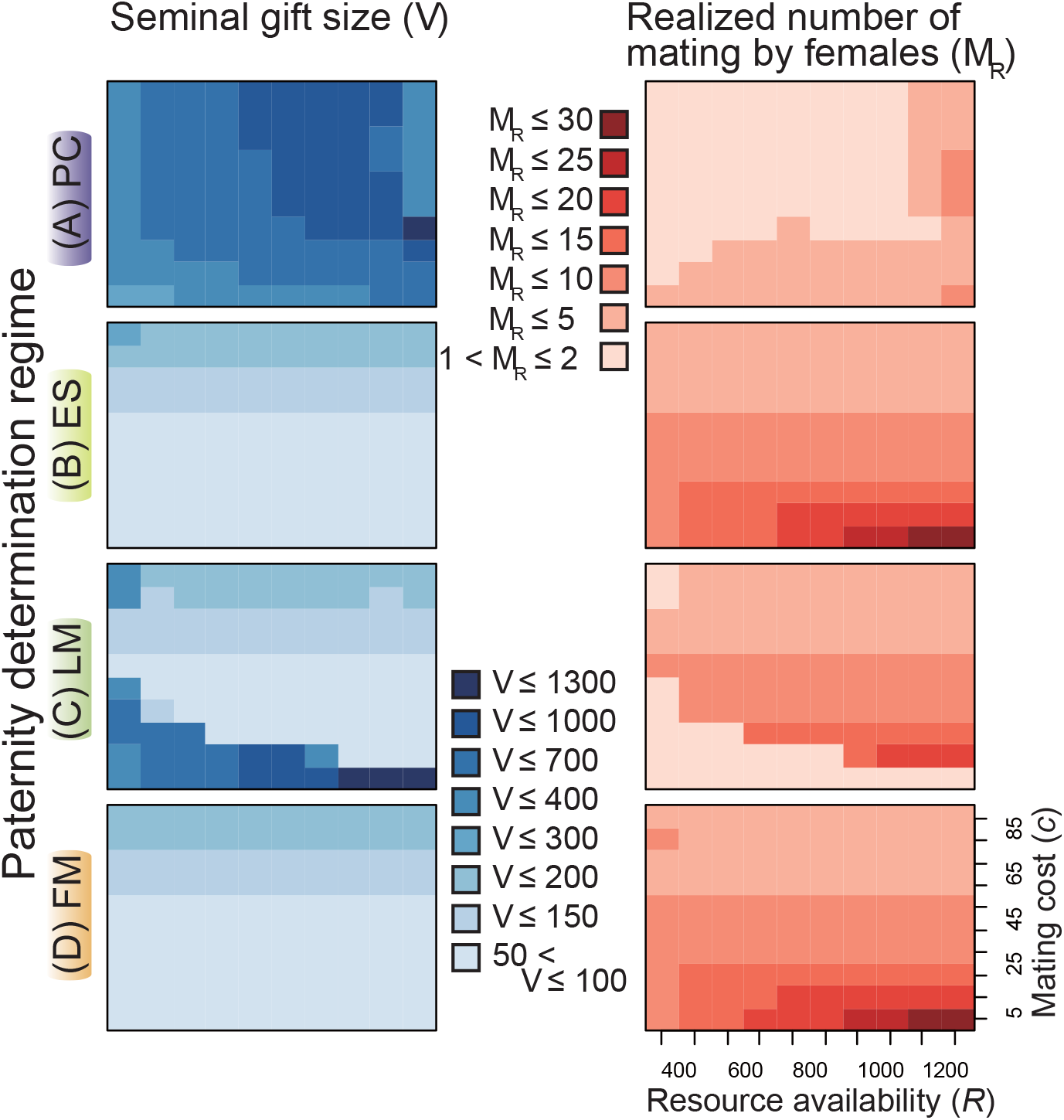
Average male seminal gift size (*V*, light blue backgrounds) and the realized number of matings by females (*M_R_*, pink backgrounds) at the 2000th generation observed in the control runs (CON) of four different paternity-determination regimes (**A**, PC; **B**, ES; **C**, LM; and **D**, FM).

Under the situations assumed in our simulation, dividing the limited resource (*R*) into small-sized gifts could increase the mating opportunities of males, while a large-sized nuptial gift could afford males two types of paternity benefits, namely: (1) siring more offspring than males who gave a smaller gift to the same mate; and (2) eliminating the probability of remating by the female as a post-copulatory guard against remating. Because the former type of benefit occurs only under the assumption of the PC regime, this can explain the observed prevalence of the “fewer large” strategy over “many small” in this regime. Accordingly, females evolve the propensity to mate multiply for these attractive, large-sized gifts when available, but males generally cannot satisfy the inflated females’ demands (Figs. 4A, 5A). Exceptions are when the mating cost is extremely large (high *c*) and the environment is eutrophic (high *R*, the upper-right corners of Figs. 4A and 5A). The former condition lowers the attractiveness of a given size of gifts, making a single mating almost optimal for females. The latter condition can shift male strategies to “many small” for seeking many virgin females, rendering their gifts more unattractive to females.

**Fig. 5.**
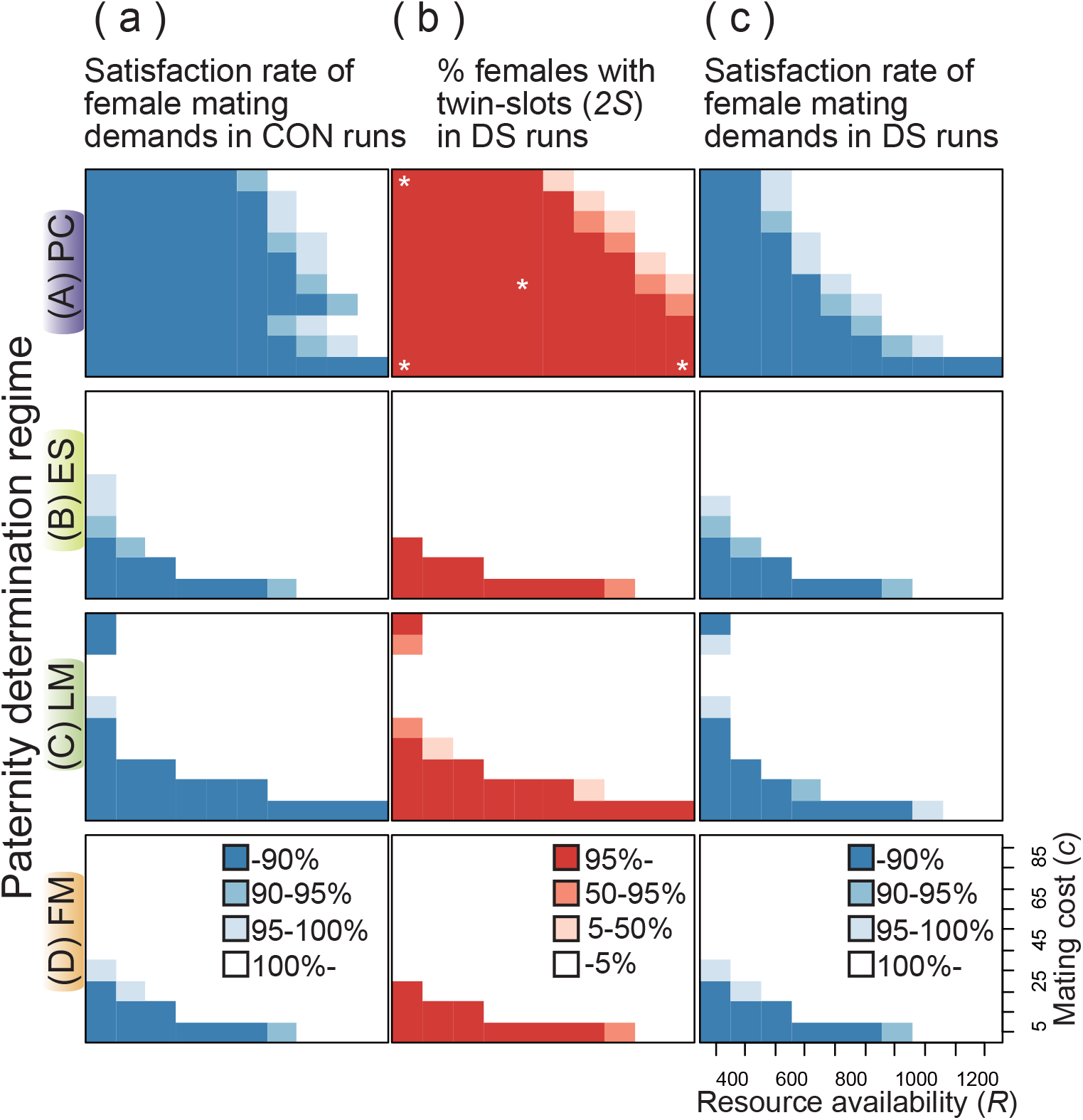
Average satisfaction rate (*M_R_*/*M_o_*) of female mating demands in CON **(a)** and DS **(c)** runs, and proportion of *2S* females **(b)** in DS runs at the 2000th generation (**A**, PC; **B**, ES; **C**, LM; and **D**, FM). The asterisks in **A-a** indicate the parameter sets examined in **Fig. 6**.

When the paternity was equally assigned to all mates of a female (ES), males generally did not evolve a large-sized nuptial gift (Fig. 4B). As discussed above, this can be attributed to the lack of paternity benefits proportional to the donated gift size in this regime. To collect these small gifts from many males, females mated more frequently than under the PC regime, especially when mating cost was low and the habitat was eutrophic (Fig. 4B). By behavioral modulation, females usually accomplished their required number of matings (Fig. 5B). An exception is the condition with very small mating costs and oligotrophic environments (the lower-left corners of Fig. 4B), where additional mating is always beneficial even for small-sized gifts.

Under the assumption that the first male monopolizes the resultant paternity (FM), males are expected to increase their opportunities to be the first mate of females by reducing the size of each gift. Interestingly, the resultant coevolutionary patterns of the nuptial gift size and the female persistence traits were almost identical between the ES and FM regimes (Figs. 4D, 5D), indicating that sexual selection works quite similarly in these two regimes.

When the last male monopolized the paternity (LM), males generally evolved larger-sized gifts compared to the FM and ES regimes (Fig. 4C). To be the last mate, we can envisage two different strategies: preparing many small gifts increases the opportunities for being the last mate by chance (as in the FM and ES regimes), while provision of a large-sized gift is advantageous because it reduces the probability of remating by the female. The latter strategy must be especially effective when females tend to seek additional mating opportunities because of low mating cost. Thus, the observed large gifts when *c* is low (the lower areas of Fig. 4C) can be a countermeasure of males for preventing frequent remating of their mates. When mating is more costly for females, it is less necessary for males to prepare large gifts, and this prudency itself reinforces the females’ reluctancy for further mating. Accordingly, for high *c* values, females satisfied their mating demands when the habitat was eutrophic (high *R*, Fig. 5C). An important exception is the extremely oligotrophic environments (*R* = 400) with extremely high cost of mating (*c* = 85–95) (the upper-left corners of Figs. 4 and 5). Under this combination of parameters, males also evolve comparatively larger gifts under the ES and FM regimes, even though it does not result in overly high mating demands of females that males cannot satisfy. Given extremely high mating costs for females, males experience a low risk of sperm competition. In addition, if males have extremely limited resources for preparing gifts, premating male-male competition should be also less severe. Accordingly, males likely shift toward providing the monandrous female as much as their resource budget allows.

In our simulation, the genetic correlation between seminal gift size ([*v1* + *v2*] / 2) and female mating propensity ([*m1* + *m2*] / 2) was negligibly low (near zero), and thus elimination of genetic covariance by shuffling paternal identity did not change the results (Supporting Material 2).

### The evolution of twin-slots and its effects on the coevolutionary processes

Figure 5 clearly shows that *2S* females successfully invaded into a population when females could not satisfy their required number of matings on average (Fig. 5). Such a condition occurs (1) when mating costs for females are extremely low and the environment is oligotrophic regardless of the paternity determination regimes, or (2) when males evolve large-sized gifts, reception of which is beneficial for females and outweighs the associated mating costs. Males evolved such an “effective” gift for making females monandrous (under the LM regime, Fig. 4C), or for siring more offspring of multiply mated females (under the PC regime, Fig. 4A).

Under these conditions, *2s* genes for making the twin slots usually increased rapidly and became fixed in the population (Fig. 3). Prevalence of *2S* females caused a coevolutionary reduction in male gift size (Fig. 3). Accordingly, females needed an increased number of matings to approach their fitness optimal, resulting in a reduction in their average fitness (Fig. 6B). When mating cost was relatively low, females could not satisfy their demands even with an increase in mating frequency by possessing two slots. However, with a higher mating cost and especially in eutrophic habitats, the smaller sized gifts, as a male counter-adaptation to twin slots, became unattractive to the females, resulting in a higher satisfaction rate (compare Fig. 5A-c with 5A-b).

**Fig. 6.**
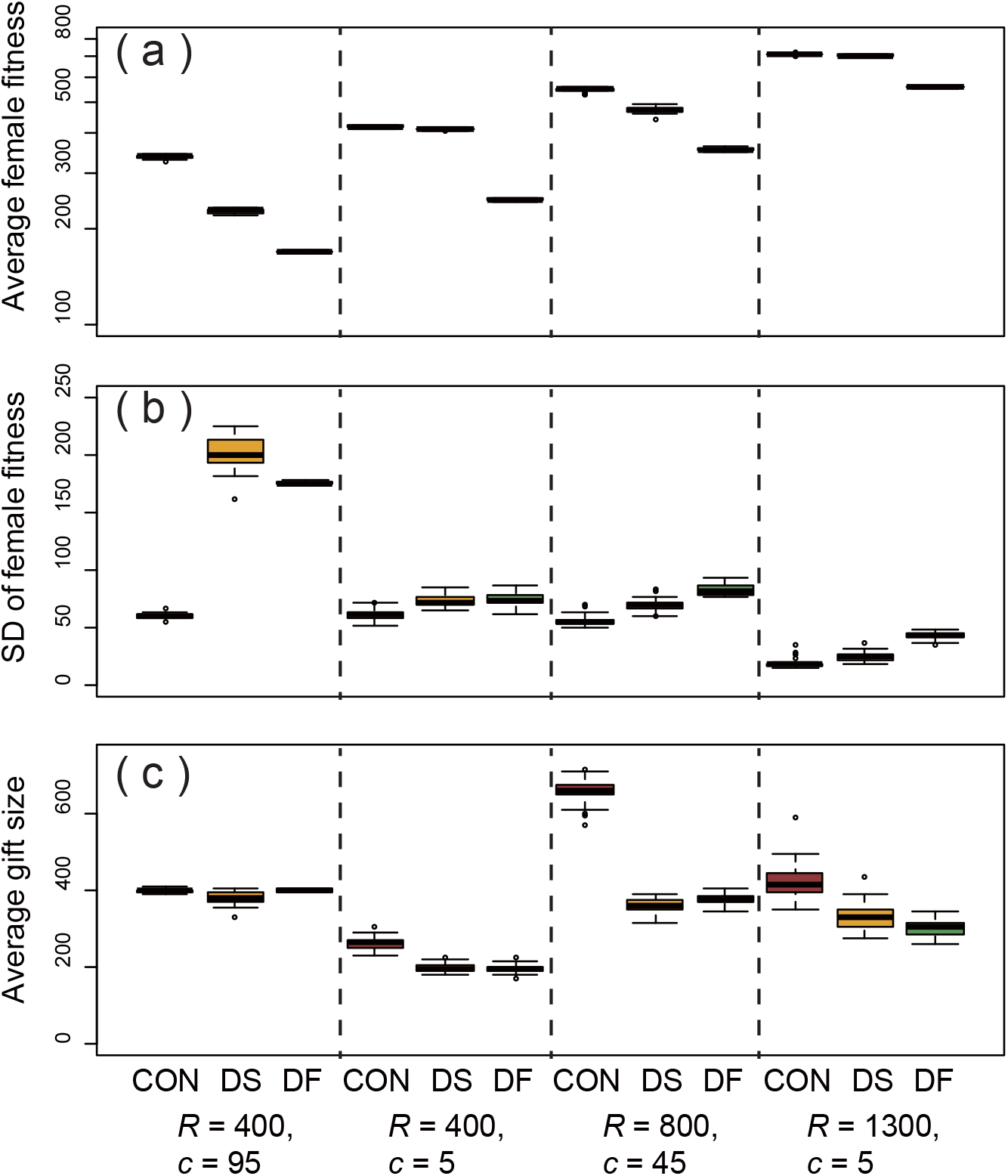
Box plots of the average female fitness **(a)**, SD of female fitness **(b)**, and average male gift size **(c)** observed at the 2000th generation under the PC regime. Results for four different parameter sets, indicated by the asterisks in **Fig. 5A-a** and the three different simulation modes (CON, control; DS, twin-slots invasion; DF, doubled number of females) are shown.

Strikingly similar patterns were observed when the number of females was doubled at the middle of simulation runs (Fig. 6A; Supporting Material 1). However, under each parameter set, the observed reduction in fitness was less prominent when *2S* females were fixed compared to when the female number was doubled (Fig. 6B), while among-individual variations in female fitness (measured as SDs) were comparative, being higher than the controls (Fig. 6C).

## 5. DISCUSSION

### Coevolutionary dynamics for trading nuptial gifts

This study demonstrated notable effects of ecological factors on the coevolutionary feedbacks among seminal gift size, female propensity for multiple mating, and twin slots for obtaining gifts. When mating costs for females are extremely low in oligotrophic environments, females compete for male derived nutrients regardless of the paternity determination regimes. This result supports the view that scarcity of other nutritive sources for female reproduction can be a favorable factor for the evolution of high dependency on male-derived gifts (Gwynne 2008; South et al. 2010). The paternity-determination regimes also show complicated interactions with these ecological factors in determining the coevolutionary feedbacks. Except for when the mating costs are negligibly low in extremely oligotrophic habitats as discussed above, both female traits, high mating rates and twin-slots, rarely evolved in the ES and FM regimes. In addition, in the LM regime, males evolved gigantic gifts almost exclusively when their donation imposed monandry of the females. A strong positive correlation between male seminal expenditure and the resultant paternity is a requisite for the evolution of effective nuptial gifts coupled with female polyandry for obtaining the gifts under a wide variety of environmental variables (Fig. 5).

There are only a few preceding studies on the coevolutionary feedbacks between the sexes in trading of ejaculate components. Like nutritious materials in nuptial gifts, sperm itself can be considered as a limited resource for both sexes (Dewsburry 1982; Nakatsuru and Kramer 1982; Olsson et al. 1997; Wedell et al. 2002; Damiens and Boivin 2006; Bro-Jørgensen 2007; Reinhardt et al. 2011; Abe 2019). Considering a situation equivalent to the PC regime of this study, a previous study examined the effects of several environmental factors on the evolutionary feedbacks between male sperm allocation strategies and female mating rate (Abe and Kamimura 2015). In the model, increased mating costs for females (*cf*, equivalent to our *c*) resulted in a lower female mating rate and an increased sperm package size, similar to the results of the present study. The study also showed that reduction in resource availability to males (low *R*) results in a monotonic reduction in the ejaculate size (Abe and Kamimura 2015). Although our present model showed similar dependency of seminal gift size (*V*) to *R*, prudently allocated gifts were also observed in extremely eutrophic environments with high female mating costs (Fig. 4A). This difference can be attributed to the different assumptions adopted: in the model of Abe and Kamimura (2015), males suffer a reduced mating rate only when females experience a shortage of sperm supply. This condition hampers the evolution of “many small” strategies when females are almost monandrous just to avoid high mating costs, while males have sufficient resources for making many seminal packages.

Bocedi and Reid (2016) also examined the effects of varying female mating costs on the coevolutionary feedbacks between male sperm traits (sperm number and sperm longevity) and female mating frequency. Under “fair raffle” sperm competition (equivalent to our PC regime), an increase in female mating costs resulted in reduced polyandry, as seen in the present study, but also in a reduction of sperm number transferred during a single mating event. The latter finding makes a striking contrast to our results in which males prepare larger gifts for less polyandrous females unless the environment is not extremely eutrophic (Fig. 4A). This can also be attributed to the different assumptions adopted for trade-off relationships: the model of Bocedi and Reid (2016) “does not explicitly consider any precopulatory male traits; overall mating frequencies are primarily determined by female mating rate,” indicating no size-number trade-offs. Instead, males increase the sperm number at a cost of reduced sperm longevity in their model, where males should invest more resources to sperm longevity when they experience a low sperm competition risk.

By modulating a “raffle loading factor,” Bocedi and Reid (2016) also found that, when the first or last males have an advantage in fertilization success (nearly equivalent to our FM or LM regimes), females tend to be more polyandrous, similar to our results. However, again, different mechanisms likely underlie these superficially similar results. Under the assumption of the number-longevity trade-off in sperm considered by Bocedi and Reid (2016), males invest less in sperm number when the first males have advantages for the sake of enhanced sperm longevity in the female sperm storage. By contrast, under the assumption of last-male sperm precedence, polyandrous mating is likely reinforced to compensate for high fertilization failure caused by reduced sperm longevity, though males provide a large number of sperm per mating (Bocedi and Reid 2016).

### Relevance to empirical studies

There is ongoing debate on the primary function of nuptial gifts for male donors, as they may function as mating efforts or paternal investments for their offspring (see Introduction). Recent studies suggest that these two hypotheses are not mutually exclusive and are difficult to discriminate, because male-derived nutrients cannot be properly allocated to the offspring of respective donors without a positive correlation between the extent of mating efforts by males and their paternity success (Lewis and South 2012). The results of the present study provide important insights for the difficulty in discriminating these two hypotheses. Nutritious nuptial gifts are effective as male mating efforts when females compete to accept (additional) mating to obtain them. The present study revealed that such effective nuptial gifts evolve almost exclusively when donation of large-sized gifts results in high paternity gains, measured as the number of offspring nourished by the female (the PC or LM regime).

Vahed (2006) explored relationships between degree of polyandry and nuptial gift size among bushcricket species. He found that nuptial gift size (spermatophylax volume) is not correlated with the degree of polyandry. However, gift size relative to body size tended to be large in species with large seminal components with possible anti-aphrodisiac effects. Thus, larger gifts may have been evolved to overcome female refractory behavior. This additional complexity was not considered in the present model.

To keep our model simple, we did not incorporate the following factors that likely affect trading of nuptial gifts: assessment of gift size by females before receiving the gift, strategic modulation of gift size by males based on mating status of mates (e.g., unmated vs. previously mated females), and costs of mating itself for males. For *Neotrogla* spp. in which copulation with the femalepositioned above the male lasts for a long period (41–73 hours in *N. curvata*, Yoshizawa et al. 2014), the last factor can be especially relevant.

We also assumed only a single reproductive output at the end of the female life (semelparity). However, in iteroparous species with multiple reproductive bouts, a male may monopolize paternity in each bout by donating a large gift, even in the cases that females are polyandrous through their life. In addition, seminal gift size can also vary interspecifically through possible macro-evolutionary trade-offs with other male and female traits: male calling frequency in bushcrickets (Del Castillo and Gwynne 2007), male weaponry in megalopteran insects (Liu et al. 2015), and male bioluminescent courtship and female flightlessness in fireflies (Lewis and Cratsley 2008; South et al. 2011; South and Lewis 2012). These complexities should be taken into account when comparing theoretical predictions with empirical data.

### Why did a “female penis” evolve in Sensitibillini

There are several hypothetical factors that have promoted the evolution of a “female penis” in *Neotrogla* (Yoshizawa et al. 2019a). In Sensitibillini, a tribe of the family Prinoglarididae with only eleven named species in the three genera known to date, females of *Afrotrogla* also possess a well-developed, presumably protrudable penis, while the homologous part forms only a small projection in the sister genus *Sensitibilla* (Lienhard 2000, 2007; Lienhard et al. 2010b). Although its function has not yet been determined, the morphology of the *Afrotrogla* female penis differs significantly from those of *Neotrogla* (Lienhard 2007, Lienhard et al. 2010a, Yoshizawa et al. 2018a). Thus, a female penis with an intromittent function is considered to have evolved twice independently in this small insect tribe (Yoshizawa et al. 2018a).

Like all other members of the tribe Sensitibillini known to date, *Neotrogla* spp. exclusively occur in oligotrophic, dry cave habitats (Lienhard et al. 2010a, Lienhard and Ferreira, 2013). In addition, they retain ancestral female-above mating positions (Fig. 2), suggesting that males do not impose high mating costs to females. Our present study demonstrates that under these conditions, that is, low *R* plus low *c*, males likely evolve large-sized gifts, for which females compete to acquire, regardless of sperm usage patterns. As in many other animals with nuptial gifts, sperm usage/storage patterns are completely unknown at present for the members of *Neotrogla*. However, the males provide considerably large gifts to polyandrous females, as evidenced by multiple emptied spermatophore capsules in the female spermatheca (Yoshizawa et al. 2014). It is also true for *Lepinotus* (Trogiidae) (Spratt 1989), females of which have only a single slot for digesting nuptial gifts (Wearing-Wilde 1995). Thus, the PC regime, in which co-occurrence of large-sized gifts and polyandry is prevalent, is the most plausible for this group of insects. Since many psocids show continuous production and deposition of eggs after sexual maturity (Söfner 1941; New 1987), the LM regime is also a candidate if females mate with a single male in each oviposition interval.

Interestingly, among Psocodea (booklice, barklice, and parasitic lice), the spermathecal duct (sperm corridor) is comparatively long and narrow in Trogiomorpha, in which formation of gigantic spermatophores is the norm, and is especially elongated and coiled in Sensitibillini (Klier 1956; Lienhard 1998, 2000; Lienhard and Ferreira 2013; Yoshizawa et al. 2019a). In the hangingfly *Hylobittacus apicalis* (Mecoptera: Bittacidae), females accept mating only while eating a nuptial gift (small arthropod prey) donated by the male. Their narrow and elongated spermathecal duct disturbs rapid sperm transfer from males. Therefore, only males who offer a large prey item are allowed to transfer enough sperm to assure their paternity (Thornhill 1976). In other words, if a spermathecal duct is wide and short, enabling rapid transfer of sperm, males may easily evolve a cheating strategy such that they pass a large number of spermatozoa with a minimal nuptial gift (see Hayashi 1999 for possible examples in megalopteran insects). Thus, it is highly plausible that the narrow and extremely long sperm corridor of Sensitibillini has a function similar to those of hangingflies, as a cryptic female choice mechanism for enabling a positive correlation between seminal gift size and transferred sperm number.

The results of the present study support the view that the evolution of twin slots can also be a crucial factor favoring the evolution of a manipulative intromittent organ in the female, as it can reinforce female-female competition for male-derived nutrients (see below). Despite the astonishing diversity in their morphology, our knowledge on the reproductive biology of the order Psocodea is quite limited. For example, although some *Neotrogla* females possess two freshly deposited spermatophores attached to the twin-slots (Yoshizawa et al. 2014), it is presently unknown whether they are derived from two different males. Future studies should also clarify the nutritional effects of spermatophores and sperm usage patterns in this unique group of insects.

### Coevolutionary feedbacks results in sex-role reversals

Our simulations demonstrated that twin slots for obtaining gifts in rapid succession are advantageous when females cannot achieve their required number of matings, and that the evolution of this persistence trait has notable effects on the coevolutionary dynamics in trading of nuptial gifts between the sexes. Little is known about the transitional process from the single-slot state, which certainly is ancestral in psocids, to the twin slots observed uniquely in Sensitibillini in the animal kingdom. The twin-slots state is estimated to have emerged from 177.5 Mya (95% confidence interval: 103.2–265.3 Mya) to 127.2 Mya (68.7.2–200.8 Mya) in a common ancestor of the extant Sensitibillini, in arid or semi-arid zones of the Pangea or Gondwana supercontinent (Yoshizawa et al. 2019b). Since the spermathecal plate of the extant Sensitibillini is a complex structure harboring not only the twin slots for seminal gifts, but also a muscle-driven mechanism to switch seminal flows between them (Yoshizawa et al. 2018b), it has likely evolved gradually from primitive precursors during its long evolutionary history of approximately 50 million years. Although we assumed that single-slot and twin-slots states can be switched by two alleles of a single locus, many associated genes must have been modified to create this evolutionarily novel structure. Future studies on the developmental processes of the spermathecal plate and their underlying genes will help us understand the difficulties of the evolution of this complex structure.

Abe and Kamimura (2015) have demonstrated that when the sex ratio is strongly biased toward females, males prepare smaller ejaculates for mating with more females. Therefore, the observed effects of the evolution of twin slots is directly comparative to those of female-biased sex ratios. However, unlike doubling the female number, the evolution of twin slots usually resulted in less pronounced reduction in female fitness with its comparatively large variance (Fig. 6). A larger variation in the female fitness means stronger sexual selection operating among them.

Male persistence traits, such as genital spines for anchoring unwilling mates or intromittent organs for traumatic insemination, can reduce the total fitness of their mates (Reinhardt et al. 2014; Tatarnick et al. 2014). Theoretical studies show that this kind of inconsistency between male and female interests can even result in a high risk of extinction, driven by the evolution of male “selfish” traits for escalated male-male competition for mates (Rankin et. al., 2011). This type of trait can also be exaggerated through arms races between the sexes so that females also develop counter-adaptations to resist or tolerate male persistence (sexually antagonistic coevolution; e.g., Brennan and Prum. 2014). There are two possible causes for the observed reduction in the average female fitness followed by the spread of *2S* females. First, twin slots themselves can directly inflate the among-female variation in cumulative amount of nutrients (*r*), and thus can reduce the average fitness under the assumption of a saturating fitness function (Fig. 1). Second, as a coevolutionary response for filling the increased number of slots, males reduce the size of each gift, resulting in an increase in the optimal number of matings for females. These lines of findings indicate that even “persistence” traits can be in the category of sex-reversed traits, driven by the evolution of effective nuptial gifts.

## REFERRENCES

Abe, J. 2019. Sperm-limited males continue to mate, but females cannot detect the male state in a parasitoid wasp. Behavioural Ecology and Sociobiology 73:52.

Abe, J., and Y. Kamimura 2015. Sperm economy between female mating frequency and male ejaculate allocation. American Naturalist 185:406–416.

Alonzo, S. H., and T. Pizzari. 2010. Male fecundity stimulation: conflict and cooperation within and between the sexes. American Naturalist 175:174–185.

Bocedi, G., and J. M. Reid. 2016. Coevolutionary feedbacks between female mating interval and male allocation to competing sperm traits can drive evolution of costly polyandry. American Naturalist 187:334–350.

Boggs, C. L. 1990. A general model of the role of male-donated nutrients in female insects’ reproduction. American Naturalist 136: 598–617.

Boggs, C. L. 1995. Male nuptial gifts: phenotypic consequences and evolutionary implications. Pages 215–242 in S. R. Leather and J. Hardie, eds. Insect Reproduction. CRC Press, Boca Raton FL

Brennan, P. L. R., and R. O. Prum. 2014. Mechanisms and evidence of genital coevolution: the roles of natural selection, mate choice, and sexual conflict. Pages 385–405 in W. R. Rice, and S. Gavrilets, eds. The genetics and biology of sexual conflict. Cold Spring harbor Laboratory Press, New York.

Chapman, T. 2001. Seminal fluid-mediated fitness traits in Drosophila. Heredity 87:511–521.

Chase, R., and K. C. Blanchard. 2006. The snail’s love-dart delivers mucus to increase paternity. Proceedings of the Royal Society B: Biological Sciences 273:1471–1475.

Damiens, D., and G. Boivin 2006. Why do sperm-depleted parasitoid males continue to mate? Behavioural Ecology 17:138–143.

Del Castillo, R. C., and D. T. Gwynne. 2007. Increase in song frequency decreases spermatophore size: correlative evidence of a macroevolutionary trade-off in katydids (Orthoptera: Tettigonidae). Evolutionary Biology 20:1028–1036.

Engqvist, L., G. Dekomien, T. Lippman, J. T. Epplen, and K. Sauer. 2007. Sperm transfer and paternity in the scorpionfly *Panorpa cognata*: large variance in traits favoured by post-copulatory episodes of sexual selection. Evolutionary Ecology 21:801–816.

Frizsche, K., and G. Arnqvist. 2013. Homage to Bateman: sex roles predict sex differences in sexual selection. Evolution 67:1926–1936.

Gwynne, D.T. 2008. Sexual conflict over nuptial gifts in insects. Annual Review of Entomology 53:83–101.

Hammerstein, P., and R. Noë. 2016. Biological trade and markets. Philosophical Transactions of the Royal Society B: Biological Sciences 371:20150101.

Hayashi, F. 1999. Rapid evacuation of spermatophore contents and male post-mating behavior in alderflies (Megaloptera: Sialidae). Entomological Science 2:49–56.

Hayashi, F., and H. Suzuki. 2003. Fireflies with or without prespermatophores: Evolutionary origins and life-history consequences. Entomological Science 6:3–10.

Kamimura, Y., and K. Yoshizawa. 2017. Sex role reversal. in J. Vonk, and T. K. Shackelford, eds. Encyclopedia of Animal Cognition and Behavior. Springer International Pub., Berlin. DOI:10.1007/978-3-319-47829-6_2012-1.

Klier, E. 1956. Zur Konstruktionsmorphologie des männlichen Geschlechtsapparates der Psocopteren. Zoologische Jahrbücher (Abteilung Anatomie) 75:207–286.

Lewis, S. M. and C. K. Cratsley. 2008. Flash signal evolution, mate choice and predation in fireflies. Annual Review of Entomology 53:293–321.

Lewis, S., and A. South. 2012. The evolution of animal nuptial gifts. Advances in the Study of Behavior 44:53–97.

Lewis, S. M., K. Vahed, J. M. Koene, L. Engqvist, L. F. Bussiere, J. C. Perry, D. Gwynne, and G. U. C. Lehmann. 2014. Emerging issues in the evolution of animal nuptial gifts. Biology Letters 10:20140336.

Lienhard, C. 1998. Psocoptères Euro–Méditerranéens. Faune de France, Paris.

Lienhard, C. 2000. A new genus of Prionoglarididae from a Namibian cave (Insecta: Psocoptera). Revue Suisse de Zoologie 107:871–882.

Lienhard, C. 2007. Description of a new African genus and a new tribe of Speleketorinae (Psocodea: ‘Psocoptera’: Prionoglarididae). Revue Suisse de Zoologie 114:441–469.

Lienhard, C., T. O. Do Carmo, and R. L. Ferreira. 2010a. A new genus of Sensitibillini from Brazilian caves (Psocodea: ‘Psocoptera’: Prionoglarididae). Revue Suisse de Zoologie 117:611–635.

Lienhard, C., O. Holuša, and G. Graffiti. 2010b. Two new cave-dwelling Prionoglarididae from Venezuela and Namibia (Psocodea: ‘Psocoptera’: Prionoglarididae). Revue Suisse de Zoologie 117:185–197.

Lienhard, C., and R. L. Ferreira. 2013 A new species of *Neotrogla* from Brazilian caves (Psocodea: ‘Psocoptera’: Prionoglarididae). Revue Suisse de Zoologie 120:3–12.

Liu, X., F. Hayashi, L. C. Lavine, and D. Yang. 2015. Is diversification in male reproductive traits driven by evolutionary trade-offs between weapons and nuptial gifts. Proceedings of the Royal Society B: Biological Sciences 282:20150247.

New, T. R. 1987. Biology of the Psocoptera. Oriental Insects 21:1–109.

Noë, R., and P. Hammerstein. 1995. Biological markets. Trends in Ecology & Evolution 10:336–339.

Perry, J. C., and L. Rowe. 2015. The evolution of sexually antagonistic phenotypes. Cold Spring Harbor perspectives in biology 7:a017558.

Rankin, D. J., U. Dieckmann, and H. Kokko. 2011. Sexual conflict and the tragedy of the commons. American Naturalist 177:780–791.

Reinhardt, K., R. Naylor, and M. T. Siva-Jothy. 2011. Male mating rate is constrained by seminal fluid availability in bedbugs, *Cimex lectularius*. PloS one 6:e22082.

Reinhardt, K., N. Anthes, and R. Lange. 2014. Copulatory wounding and traumatic insemination. Pages 115–139 in W. R. Rice and S. Gavrilets, eds. The genetics and biology of sexual conflict. Cold Spring harbor Laboratory Press, New York.

Sauer, K. P., T. Lubjuhn, J. Sindern, H. Kullmann, J. Kurtz, C. Epplen, and J. T. Epplen 1998. Mating system and sexual selection in the scorpionfly *Panorpa vulgaris* (Mecoptera: Panorpidae). Naturwissenschaften 85:219–228.

Simmons, L. W., and G. A. Parker. 1989. Nuptial feeding in insects: mating effort versus paternal investment. Ethology 81:332–343.

Söfner, L. 1941. Zur Entwicklungsbiologie und Ökologie der einheimischen Psocopterenarten *Ectopsocus meridionalis* Ribaga 1904 und *Ectopsocus briggsi* McLachlan 1899. Zoologische Jahrbücher (Abteilung Systematik) 74:323–360.

South, A., and S. M. Lewis. 2012. Determinants of reproductive success across sequential episodes of sexual selection in a firefly. Proceedings of the Royal Society B: Biological Sciences 279:3201–3208.

South, A, K. Stanger-Hall, M.-L. Jeng, and S. M. Lewis. 2011. Correlated evolution of female neoteny and flightlessness with male spermatophore production in fireflies (Coleopteta: Lampyridae). Evolution 65:1099–1113.

Spratt, E. C. 1989. The incidence of spermatophores and the possible significance of their formation in *Lepinotus patruelis* Pearman (Psocoptera: Trogiidae). Entomologist’s Gazette 40:235–239.

Tatarnic, N. J., G. Cassis, and M. T. Siva-Jothy. 2014. Traumatic insemination in terrestrial arthropods. Annual Review of Entomology 59:245–261.

Vahed, K. 1998. The function of nuptial feeding in insects: a review of empirical studies. Biological Reviews 73:43–78.

Vahed, K. 2006. Larger ejaculate volumes are associated with a lower degree of polyandry across bushcricket taxa. Proceedings of the Royal Society B: Biological Sciences 273:2387–2394.

Wearing-Wilde, J. 1995. The sclerotized spermatophore of the barklouse *Lepinotus patruelis*. Tissue and Cell 27:447–456.

Wearing-Wilde, J. 1996. Mate choice and competition in the barklouse *Lepinotus patruelis* (Psocoptera: Trogiidae): The effect of diet quality and sex ratio. Journal Of Insect Behavior. 9:599–612.

Wedell, N., M. J. Gage, and G. A. Parker. 2002. Sperm competition, male prudence and sperm-limited females. Trends in Ecology and Evolution 17:313–320.

Yoshizawa, K., R. L. Ferriera, Y. Kamimura, and C. Lienhard. 2014. Female penis, male vagina, and their correlated evolution in a cave insect. Current Biology 24:1006–1010.

Yoshizawa, K., R. L. Ferreira, I. Yao, C. Lienhard, and Y. Kamimura. 2018a. Independent origins of female penis and its coevolution with male vagina in cave insects (Psocodea: Prionoglarididae). Biology Letters 14:20180533.

Yoshizawa, K., Y. Kamimura, C. Lienhard, R. L. Ferreira, and A. Blanke 2018b. A biological switching valve evolved in the female of a sex-role reversed cave insect to receive multiple seminal packages. eLife 7:e39563.

Yoshizawa, K., R. L. Ferreira, C. Lienhard, and Y. Kamimura. 2019a. Why did a female penis evolve in a small group of cave insects? BioEssays 41:1900005.

Yoshizawa, K., C. Lienhard, I. Yao, and R. L. Ferreira. 2019b. Cave insects with sex-reversed genitalia had their most recent common ancestor in West Gondwana (Psocodea: Prinoglarididae: Speleketorinae). Entomological Science. 22:334–338.

